# Deleterious mutation accumulation in *Arabidopsis thaliana* pollen genes: a role for a recent relaxation of selection

**DOI:** 10.1101/016626

**Authors:** MC Harrison, EB Mallon, D Twell, RL Hammond

## Abstract

In many studies sex related genes have been found to evolve rapidly. We therefore expect plant pollen genes to evolve faster than sporophytic genes. In addition, pollen genes are expressed as haploids which can itself facilitate rapid evolution because recessive advantageous and deleterious alleles are not masked by dominant alleles. However, this mechanism is less straightforward to apply in the model plant species *Arabidopsis thaliana*. For 1 million years *A.thaliana* has been self-compatible, a life history switch that has caused: a reduction in pollen competition, increased homozygosity and a dilution of masking in diploid expressed, sporophytic genes. In this study we have investigated the relative strength of selection on pollen genes compared to sporophytic genes in *A. thaliana*. We present two major findings: 1) before becoming self-compatible positive selection was stronger on pollen genes than sporophytic genes for *A. thaliana*; 2) current polymorphism data indicate selection is weaker on pollen genes compared to sporophytic genes. These results indicate that since *A. thaliana* has become self-compatible, selection on pollen genes has become more relaxed. This has led to higher polymorphism levels and a higher build-up of deleterious mutations in pollen genes compared to sporophytic genes.

## Introduction

A faster evolution of reproductive genes compared to somatic genes has been documented for a wide range of taxa, including primates, rodents, mollusks, insects and fungi (Turner and Hoekstra, 2008; Swanson and Vacquier, 2002). The faster evolution is often observable in a higher number of non-synonymous nucleotide substitutions (base changes which alter the amino acid sequence of a protein) within the coding regions of orthologues. In most cases stronger positive selection is described as the mechanism driving the divergence of these genes, generally due to some form of sexual selection like cryptic female choice or sperm competition.

Two studies on the strength of selection on reproductive and non-reproductive genes in *Arabidopsis thaliana* presented somewhat conflicting findings (Szövényi *et al.*, 2013; Gossmann *et al.*, 2013). Szövényi *et al.* (2013) showed that the rate of protein evolution, measured in terms of dN/dS (ratio of non-synonymous to synonymous per site substitution rates), between *Arabidopsis thaliana* and *A. lyrata* of pollen-specific genes was significantly higher than for sporophyte-specific genes (Szövényi *et al.*, 2013). The detection of higher intra-specific polymorphism levels within pollen genes was compatible with relaxed purifying selection on pollen genes. This is because stronger positive selection, that could have caused the higher divergence rates, would have reduced intra-specific polymorphism levels. High tissue specificity and higher expression noise compared to sporophytic genes were considered the likely causes of relaxed selection on pollen genes. As pointed out in a further study, which focused on the comparison of genes with male biased or female biased expression (Gossmann *et al.*, 2013), inter-specific divergence and currently existing intra-specific polymorphisms likely arose under different selection regimes for *A. thaliana*. The divergence of *A. thaliana* from its closest relative *A. lyrata* happened largely during a period of outcrossing, since speciation occurred approximately 13 million years ago (Beilstein *et al.*, 2010), whereas *A. thaliana* became self-compatible only roughly one million years ago (Tang *et al.*, 2007). Divergence patterns for *A. thaliana* should therefore be similar to outcrossing species and reveal stronger selection on pollen genes. Existing, intra-specific polymorphisms, on the other hand, are expected to be influenced by high selfing rates in *A. thaliana* populations that have led to high levels of homozygosity across the whole genome (Nordborg, 2000; Wright *et al.*, 2008; Platt *et al.*, 2010). The outcome is a reduction in the masking of deleterious alleles in diploid sporophyte stages (because of high homozy-gosity) compared to the haploid gametophyte stage. Furthermore, selfing will result in fewer genotypes competing for fertilization so lowering the magnitude of pollen competition and reducing the strength of selection acting on pollen (Charlesworth and Charlesworth, 1992).

Gossmann *et al.* (2013) found protein divergence (dN/dS) to be higher for female biased genes compared to both male genes and 476 random, non-reproductive genes sampled from the *A. thaliana* genome. However, pollen genes did not differ from the non-reproductive genes in terms of dN/dS. Despite using a larger number of accessions to measure polymorphism than in the Szövényi *et al.* study (80 compared to 19), Gossmann *et al.* did not detect any difference in nucleotide diversity between the non-reproductive genes and pollen-specific genes in general, although nucleotide diversity was significantly lower for sperm cell-specific genes (Gossmann *et al.*, 2013). When comparing polymorphism to divergence data with a modified version of the McDonald-Kreitman test (McDonald and Kreitman 1991, Distribution of Fitness Effects Software, DoFE; Eyre-Walker and Keightley 2009) a higher proportion of non-synonymous sites were found to be under purifying and adaptive selection for pollen genes compared to both female biased and non-reproductive genes.

The aim of our study was to attempt to resolve these apparently conflicting results for *A. thaliana* and to address the following questions. Are pollen proteins really more divergent than sporophyte proteins? If so, is this due to more relaxed purifying selection or increased positive selection on pollen genes? Have patterns of selection changed for *A. thaliana* since it became self-compatible? In a first step we estimated the protein divergence of 1,552 pollen and 5,494 sporophytic genes to both *A. lyrata* and *Capsella rubella* in terms of interspecific dN/dS. This larger gene set, combined with a larger number of accessions than both previous studies (269 compared to 80 and 19), increased the power to detect sites under positive and negative selection within the two groups of genes when conducting a DoFE analysis. As the polymorphism and divergence data likely reflect periods of differing selection regimes (divergence under self incompatibility, polymorphism under self compatibility) we additionally detected sites under positive selection using a site model of the Phylogenetic Analysis by Maximum Likelihood software (PAML 4.6; Yang 2007), which does not require polymorphism data and detects sites under positive selection by allowing dN/dS to vary within genes. In a second step, to investigate more recent selection patterns, we analyzed intra-specific polymorphism levels within each group of genes. Lower diversity, measured here via non-synonymous Watterson’s *θ* and nucleotide diversity (*π*), would be expected for pollen genes compared to sporophyte genes in the case of stronger selection (Nielsen, 2005). In a further test we also compared existing levels of putative deleterious alleles (premature stop codons and frameshift mutations) between pollen genes and sporophyte genes. In each of these analyses we controlled for differences in genomic factors (expression level, GC content, codon bias, gene density, gene length and average intron length) between the pollen and sporophyte-specific genes which were correlated with the divergence, polymorphism and deleterious allele measurements.

## Materials and Methods

### Genomic data

Publicly available variation data were obtained for 269 inbred strains of *A. thaliana*. Beside the reference genome of the Columbia strain (Col-0), which was released in 2000 (*Arabidopsis*, Genome Initiative), 250 were obtained from the 1001 genomes data center (http://1001genomes.org/datacenter/; accessed September 2013), 170 of which were sequenced by the Salk Institute (Schmitz *et al.*, 2013) and 80 at the Max Planck Institute, Tübingen (Cao *et al.*, 2011). A further 18 were downloaded from the 19 genomes project (http://mus.well.ox.ac.uk/; accessed September 2013; Gan *et al.* 2011). These 269 files contained information on SNPs and indels recorded for separate inbred strains compared to the reference genome. A quality filter was applied to all files, in order to retain only SNPs and indels with a phred score of at least 25. For further analyses, gene sequences were created for each of these strains based on coding sequence information contained in the TAIR10 gff3 file.

### Expression data

Normalized microarray data, covering 20,839 genes specific to different developmental stages and tissues of *A. thaliana* (table 10), were obtained from Borg *et al.* (2011). The expression data consisted of 7 pollen and 10 sporophyte data sets (table 10). Four of the pollen data sets represented expression patterns of the pollen developmental stages, uninucleate, bicellular, tricellular and mature pollen grain, one contained expression data of sperm cells and the remaining two were pollen tube data sets. There was a strong, significant correlation between the two pollen tube data sets (*ρ* = 0.976; p *<* 2.2×10^−16^; Spearman’s rank correlation), so both were combined and the highest expression value of the two sets was used for each gene. Each of the 10 sporophyte data sets contained expression data for specific sporophytic tissues (table 10).

Each expression data point consisted of a normalized expression level (ranging from 0 to around 20,000, scalable and linear across all data points and data sets) and a presence score ranging from 0 to 1 based on its reliability of detection across repeats, as calculated by the MAS5.0 algorithm (Borg *et al.*, 2011). In our analyses expression levels were conservatively considered as present if they had a presence score of at least 0.9, while all other values were regarded as zero expression.

Genes were classed as either pollen or sporophyte-specific genes, if expression was reliably detectable in only pollen or only sporophyte tissues or developmental stages. The highest expression value across all tissues or developmental stages was used to define the expression level of a particular gene. The highest value was used since this best represents the genes’ most important effect on the phenotype. We also consider tissue specificity of expression to fully explain a gene’s expression profile.

### Detecting signatures of selection

#### Evolutionary Rates

To estimate evolutionary rates of genes, dN/dS ratios (ratio of non-synonymous to synonymous substitution rates relative to the number of corresponding non-synonymous and synonymous sites) were calculated for all orthologous genes between pairs of the three species *A. thaliana*, *A. lyrata* and *Capsella rubella* using the codeml program within the PAML package (Yang, 2007). The protocol described in Szövényi *et al.* (2013) was followed. Orthologues were found by performing reciprocal blastp searches (Altschul *et al.*, 1997) between proteomes and retaining protein pairs with mutual best hits showing at least 30% identity along 150 aligned amino acids (Rost, 1999). Orthologous protein sequences were aligned with MUSCLE (Edgar, 2004) at default settings and mRNA alignments were performed based on these protein alignments with pal2nal (Suyama *et al.*, 2006). The codeml program was run with runmode -2, model 2 and ’NSsites’ set to 0. In most results we report divergence (dN/dS) between *A. thaliana* and *A. lyrata* unless otherwise stated.

In order to detect genes that contain codon sites under positive selection, we performed a likelihood-ratio test (LRT) between models 7 (null hypothesis; dN/dS limited between 0 and 1) and 8 (alternative hypothesis; additional parameter allows dN/dS *>* 1) by using runmode 0, model 0 and setting ’NSsites’ to 7 & 8. An LRT statistic (twice the difference in log-likelihood between the two models) greater than 9.210 indicated a highly significant difference (p *<* 0.01; LRT *>* 5.991: p *<* 0.05) between the two models suggesting the existence of sites under positive selection within the tested gene (Anisimova *et al.*, 2003; Yang, 2007). These tests were carried out on multi-species alignments containing orthologues from *A. thaliana*, *A. lyrata* and *C. rubella* that were contained in each of the three orthologue lists described before. Alignments were carried out in the same manner as described above for pairs of sequences.

Levels of purifying and positive selection were estimated with the Distribution of Fitness Effects Software (DoFE 3.0) using the Eyre-Walker and Keightley (2009) method. For the input files synonymous and non-synonymous site spectra were obtained using the Pegas package (Paradis, 2010) in R (version 3.2.0; R Core Team 2012). Four-fold sites were used to represent synonymous positions and zero-fold degenerate sites to represent nonsynonymous positions. Four-fold and zero-fold sites were calculated with perl scripts; any codons containing more than one SNP were removed from the analysis. We randomly sampled 20 alleles at each site without replacement using the perl module ’shuffle’. Ten analyses were carried out for each gene group, each time randomly sampling 50 genes without replacement. Values were summed across all genes. The results were checked for convergence as advised in the user manual and repeated with new samples if any overall trends were observed. The results of all 10 samples were combined and presented in this study.

#### Intra-specific polymorphism

Nucleotide diversity (*π*) and Watterson’s *θ* were calculated for non-synonymous sites using the R package PopGenome (version 2.1.6; Pfeifer *et al.* 2014). The diversity.stats() command was implemented and the subsites option was set to ”nonsyn”. Both values were subsequently divided by the number of sites.

#### Putatively deleterious alleles

To quantify the frequency of deleterious mutations for each gene, the occurrence of premature stop codons and frameshifts was calculated for each gene locus across all 268 strains compared to the reference genome. Stop codons were recorded as the number of unique alternative alleles occurring within the 269 strains as a result of a premature stop codon. Frameshifts were calculated as a proportion of the strains containing a frameshift mutation for a particular gene. All analyses of coding regions were based on the representative splice models of the *A. thaliana* genes (TAIR10 genome release, www.arabidopsis.org).

#### Statistical analyses

All analyses were performed in R (version 3.2.0; R Core Team 2012). To measure statistical difference between groups we utilized the non-parametric Mann Whitney U test (wilcox.test() function). In case of multiple testing, all p-values with corrected with the Bonferroni method using the function p.adjust(). For correlations either the Spearman rank test (rcorr() function of Hmisc package; version 3.16-0; Jr and others 2015) or Spearman rank partial correlation (pcor.test() function; ppcor package; version 1.0; Kim 2012) was carried out.

Six genomic parameters were investigated as possible predictors of dN/dS, polymorphism levels and frequency of deleterious mutations. These were expression level, GC-content, codon bias variance, gene density, average gene length and average intron length. Expression level is described above in the section ”Expression data”. Average gene length and average intron length were calculated using custom made scripts which extracted information from the genomic gff file. GC content was calculated with a downloaded Perl script, which was originally written by Dr. Xiaodong Bai (http://www.oardc.ohio-state.edu/tomato/HCS806/GC-script.txt). RSCU (relative synonymous codon usage) was used to measure codon bias. It was calculated for each codon of each locus with the R package ’seqinr’ (uco() function; version 3.1-3; Perriere 2014). As the mean value per gene varied very little between loci but varied by site within genes, we used RSCU variance as a measure for codon bias. Gene density was calculated with custom Perl and R scripts by counting the number of genes within each block of 100kb along each chromosome. Gene densities were then attributed to each gene depending on the 100kb window, in which they were situated.

As most of the genomic parameters investigated here (gene expression, GC-content, codon bias variance, gene density, average gene length and average intron length) generally differed between groups of genes (see Results), it was important to control for their possible influence on divergence, polymorphism and frequencies of deleterious mutations. The 6 parameters were also inter-correlated, so we decided to implement principle component regression analyses (pcr() command, pls package, version 2.4-3; Mevik and Wehrens 2007) in order to combine these parameters into independent predictors of the variation in the investigated dependent variable (e.g. dN/dS) as described by Drummond *et al.* (2006). All variables, including the dependent variable, were log transformed (0.0001 was added to gene length and average intron length due to zero values). A jack knife test (jack.test()) was subsequently performed on each set of principal component regression results to test if the contribution of each predictor was significant. Non-significant predictors were then removed and the analyses were repeated. The principle component (PC), which explained the highest amount of variation in the dependent variable, was then used to represent the genomic predictors in an ANCOVA (e.g. lm(log(dN/dS) ~ PC1 * ploidy)) with life-stage as the binary co-variate.

## Results

### Life-stage limited genes

Within the total data set, containing 20,839 genes, 4,304 (20.7%) had no reliably detectable expression (score *<* 0.9; see methods) in any of the analysed tissues and were removed from the analysis. Of the remaining 16,535 genes, 1,552 genes (9.4%) were expressed only in pollen and a further 5,494 (33.2%) were limited to sporophytic tissues (referred to as pollen-specific genes and sporophyte-specific genes in this study). The pollen-specific and sporophyte-specific genes were randomly distributed among the five chromosomes (table 1), and their distributions within the chromosomes did not differ significantly from each other (table 2).

**Table 1:** Chi squared test of the distribution of pollen and sporophyte limited genes among the five nuclear *A. thaliana* chromosomes. Degrees of freedom: 4.

| Chromosome | All genes | Pollen | Sporophyte |
| --- | --- | --- | --- |
| 1 | 4,348 | 392 | 1,495 |
| 2 | 2,522 | 251 | 862 |
| 3 | 3,326 | 340 | 1,049 |
| 4 | 2,451 | 214 | 839 |
| 5 | 3,888 | 355 | 1,249 |
| $\Sigma$ | 16,535 | 1,552 | 5,494 |
| $\chi^2$ | | 5.367 | 7.456 |
| p |  | 0.252 | 0.136 |

**Table 2:** Comparison of chromosomal positions of pollen and sporophyte genes. Mann Whitney U test.

| Chromosome | W | p |
| --- | --- | --- |
| 1 | $2.79 \times 10^5$ | 0.137 |
| 2 | $1.00 \times 10^5$ | 0.071 |
| 3 | $1.72 \times 10^5$ | 0.315 |
| 4 | $8.54 \times 10^4$ | 0.267 |
| 5 | $2.31 \times 10^5$ | 0.241 |

Expression level was roughly twice as high within pollen-specific genes (median: 1,236.1) compared to sporophyte-specific genes (median: 654.7; W = 5.5 x 10^6^; p = 1.2 x 10^−63^; table 3). GC-content was significantly higher within sporophyte-specific genes (median: 44.6%) than in pollen-specific genes (median: 43.8%; W = 3.4 x 10^6^; p = 1.0 x 10^−19^; table 3). Sporophyte-specific genes were significantly longer and contained significantly longer introns than pollen-specific genes (table 3). Gene density was slightly but significantly higher in pollen-specific genes; codon bias variance did not differ significantly (table 3).

**Table 3:** Differences in 6 genomic variables between pollen-specific and sporophyte-specific genes. Values are means standard error of the mean; significance was tested with Mann Whitney U test ± p-values are Bonferroni corrected for multiple testing.

| Genomic variable | Pollen-specific genes |  |  | Sporophyte-specific genes |  |  | p |
| --- | --- | --- | --- | --- | --- | --- | --- |
| Expression level | 2,562.30 | ± 86.49 | > | 1,256.21 | ± 23.80 | | $1.2 \times 10^{-63}$ |
| GC content (%) | 44.20 | ± 0.08 | < | 45.08 | ± 0.04 | | $1.0 \times 10^{-19}$ |
| Codon bias variance | 0.46 | ± 0.01 | = | 0.43 | ± 0.00 | | not significant |
| gene length | 1,570.30 | ± 24.41 | < | 1,634.39 | ± 11.62 | | $2.3 \times 10^{-4}$ |
| average intron length | 124.44 | ± 3.23 | < | 160.08 | ± 2.49 | | $8.6 \times 10^{-10}$ |
| gene density (per 100kb) | 29.99 | ± 0.12 | > | 29.57 | ± 0.07 | | $1.5 \times 10^{-3}$ |

### Pollen-specific proteins evolve at a faster rate than sporophyte-specific proteins

The rate of evolution of *Arabidopsis thaliana* proteins from *Arabidopsis lyrata* orthologues was estimated using interspecific dN/dS. Of the 13,518 genes for which 1-to-1 orthology could be detected and dN/dS could be reliably analysed, 1144 genes were pollen-specific and 4395 were sporophyte-sepecific. Protein divergence was significantly higher for pollen-specific genes than sporophyte-specific genes (p = 4.3 x 10^−24^; table 6, fig. 1(c)). This was mainly due to a significant difference in the non-synonymous substitution rate for which the median was 30.8% higher in pollen specific genes (dN; p = 2.4 x 10^−27^; fig. 1(a)). Synonymous divergence (dS) was only 3.7% higher in pollen-specific genes and the difference was less significant (p = 1.6 x 10^−4^; fig. 1(b)).

**Figure 1:**
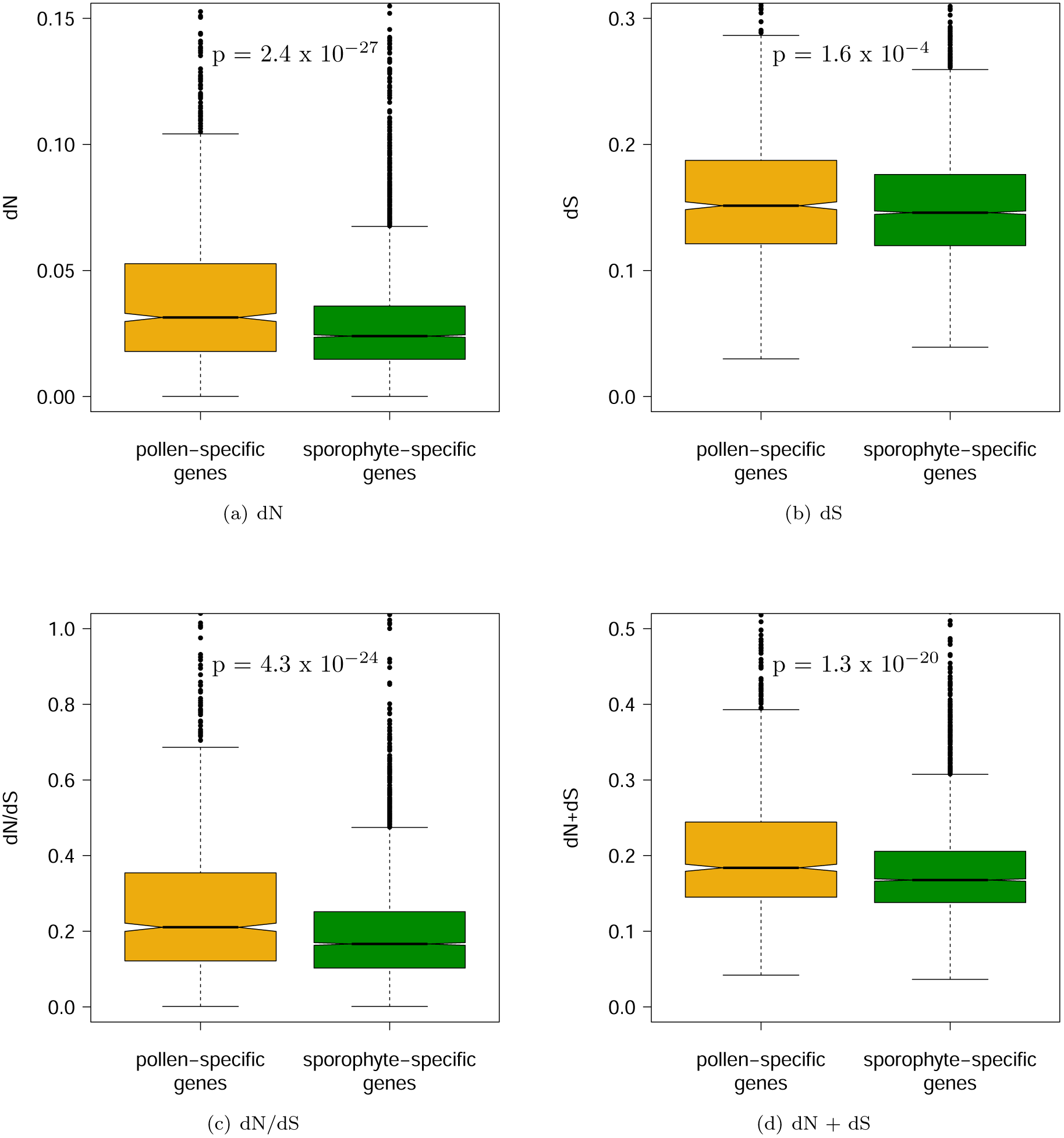
Non-synonymous (dN; a), synonymous (dS; b), dN/dS (c) and total nucleotide substitution rate (dN + dS; d) within pollen-specific and sporophyte-specific genes. Significance tested with Mann Whitney U test.

Both expression level (*ρ* = -0.232; p = 5.6 x 10^−169^) and GC-content (*ρ* = -0.145; p = 4.3 x 10^−64^) were significantly negatively correlated with dN/dS while controlling for other factors (codon bias variance, gene length, average intron length and gene density; table 4). Codon bias variance and gene length correlated weakly and negatively with dN/dS, while average intron length and gene density showed minimal correlation (table 4).

**Table 4:**
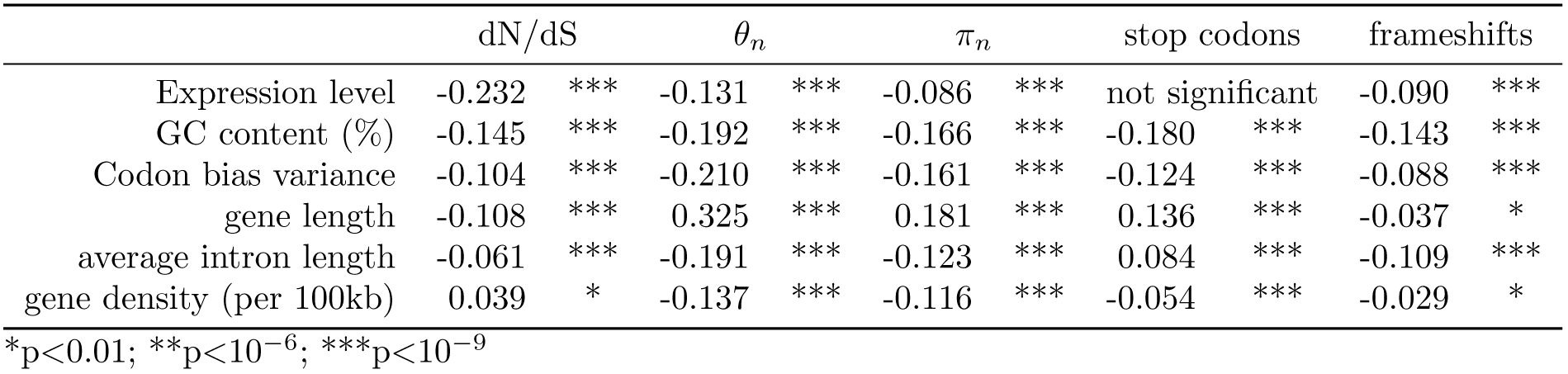
Partial correlations of 6 genomic variables with dN/dS, *θ_n_*, *π_n_*, frequency of premature stop codons and frameshift mutations. Spearman rank correlations controlling for remaining 5 variables; p-values are Bonferroni corrected for multiple testing.

In order to determine how the life-stage (pollen or sporophytic tissue), to which the expression of a gene is limited, may be contributing to the measured difference in dN/dS, it was important to control for the six previously mentioned genomic variables (expression level, GC-content, codon bias variance, gene length, average intron length and gene density). This was important since five of the six genomic variables differed significantly between pollen and sporophyte-specific genes (table 3) and all six were significantly correlated to dN/dS (table 4). A principal component regression was conducted to allow us to condense these predictors of dN/dS into independent variables. We first included all 6 predictors in the principal component regression model, and they explained 9.10% of dN/dS variation. Principal component (PC) 2 explained the largest amount of variation at 6.15%. A jack knife test on this PC revealed significant p-values (*<* 0.05) only for expression, GC content and codon bias variance. After removal of the non-significant predictors (gene length, average intron length and gene density) codon bias variance was also no longer significant. The first PC of a model containing expression and GC content as the predictors of dN/dS had an explanation value of 7.15% (total 7.24%). This first PC was used as the continuous variable in an ANCOVA with dN/dS as the dependent variable and life-stage as the binary co-variable. The pollen regression line was higher than for the sporophyte genes for the majority of the PC1 range (fig. 2). As the slopes differed significantly (p = 4.4 x 10^−4^), we measured the difference in dN/dS between pollen and sporophyte genes within 5 equal bins along the PC1 axis. In all five quantiles dN/dS was higher within pollen genes than within sporophyte-specific genes (highly significant in the first three, marginally significant in the fourth after correction and non-significant in the fifth quantile; table 5).

**Figure 2:**
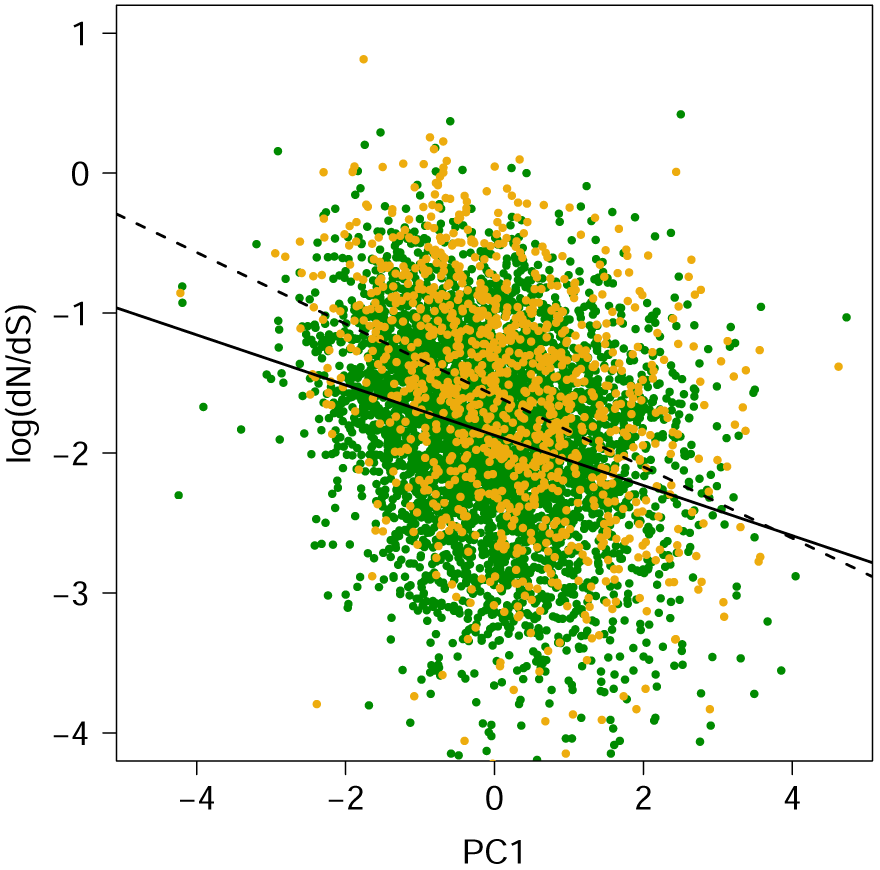
ANCOVA analysis of dN/dS within pollen-specific (yellow points and dashed line) sporophyte-specific genes (green points and solid line) with PC1 (expression and GC content) as the continuous variable.

**Table 5:** dN/dS within 5 equal bins along the PC1 axis. Shown are medians (means).

|  | < 20% |  | 20% - 40% |  | 40% - 60% |  | 60% - 80% |  | > 80% |  |
| --- | --- | --- | --- | --- | --- | --- | --- | --- | --- | --- |
| Pollen | 0.269 | (0.347) | 0.249 | (0.346) | 0.210 | (0.276) | 0.173 | (0.214) | 0.144 | (0.198) |
| Sporophyte | 0.220 | (0.247) | 0.173 | (0.199) | 0.160 | (0.183) | 0.146 | (0.192) | 0.132 | (0.160) |
| p | 3.7 x 10<sup>-5</sup> |  | 1.4 x 10<sup>-9</sup> |  | 1.6 x 10<sup>-6</sup> |  | 0.050 |  | non-significant |  |

### Sporophyte-specific genes contain a higher number of sites under purifying selection

We investigated whether the higher divergence of pollen-specific proteins compared to sporophyte-specific proteins was restricted to *Arabidopis*, and possibly fueled by selection in either *A. thaliana* or *A. lyrata*, by investigating the protein divergence of both from *Capsella rubella*. Divergence was significantly higher for pollen-specific proteins in all three comparisons (table 6). Between branches only one comparison of divergence values differed significantly for sporophyte-specific proteins: *A. thaliana*-*A. lyrata* dN/dS *> A. lyrata*-*C. rubella* dN/dS (Bonferroni corrected p-value: 0.046); all other differences between branches were non-significant.

**Table 6:** dN/dS between *A. thaliana*, *A. lyrata* and *C. rubella*. Values are means (and medians); significance was tested with Mann Whitney U test; p-values are Bonferroni corrected for multiple testing.

|  | Pollen | Sporophyte | p value |
| --- | --- | --- | --- |
| *A. thaliana* vs. *A. lyrata* | 0.2689 (0.2106) | 0.1963 (0.1664) | 4.3 x 10<sup>-24</sup> |
| *A. thaliana* vs. *C. rubella* | 0.2409 (0.2036) | 0.1801 (0.1567) | 8.8 x 10<sup>-22</sup> |
| *A. lyrata* vs. *C. rubella* | 0.2370 (0.1945) | 0.1818 (0.1568) | 1.3 x 10<sup>-15</sup> |

A higher dN/dS value, which is still lower than 1, generally indicates weaker purifying selection (Yang and Bielawski, 2000). Only 41 out of 13,518 genes had a dN/dS value greater than 1 and 65.1% of genes had a dN/dS less than 0.2. However, gene-wide estimates of dN/dS can be inflated by a few codon sites under positive selection (*>* 1) even if purifying selection is otherwise prevalent. In order to test whether the higher dN/dS within pollen genes was being driven by relaxed purifying selection or increased positive selection we analysed the distribution of fitness effects of new mutations using the DoFE software (DoFE 3.0; Eyre-Walker and Keightley 2009). The analyses were repeated 10 times on random samples of 20 alleles and 50 genes from each group. The distribution of new deleterious mutations showed that a smaller fraction of non-synonymous mutations were strongly deleterious (N_*e*_s *>* 10; N_*e*_: effective population size, s: selection coefficient) within pollen-specific genes (mean 42.9%) compared to sporophyte genes (47.9%; Fig. 3). Also, a higher proportion of mutations in pollen genes were effectively neutral (N_*e*_s *<* 1; 51.0%) compared to sporophyte genes (45.7%). This indicates weaker purifying selection within the pollen-specific genes (Eyre-Walker and Keightley, 2009) and suggests the higher dN/dS rates in pollen genes may be caused by an accumulation of slightly deleterious mutations due to random drift.

**Figure 3:**
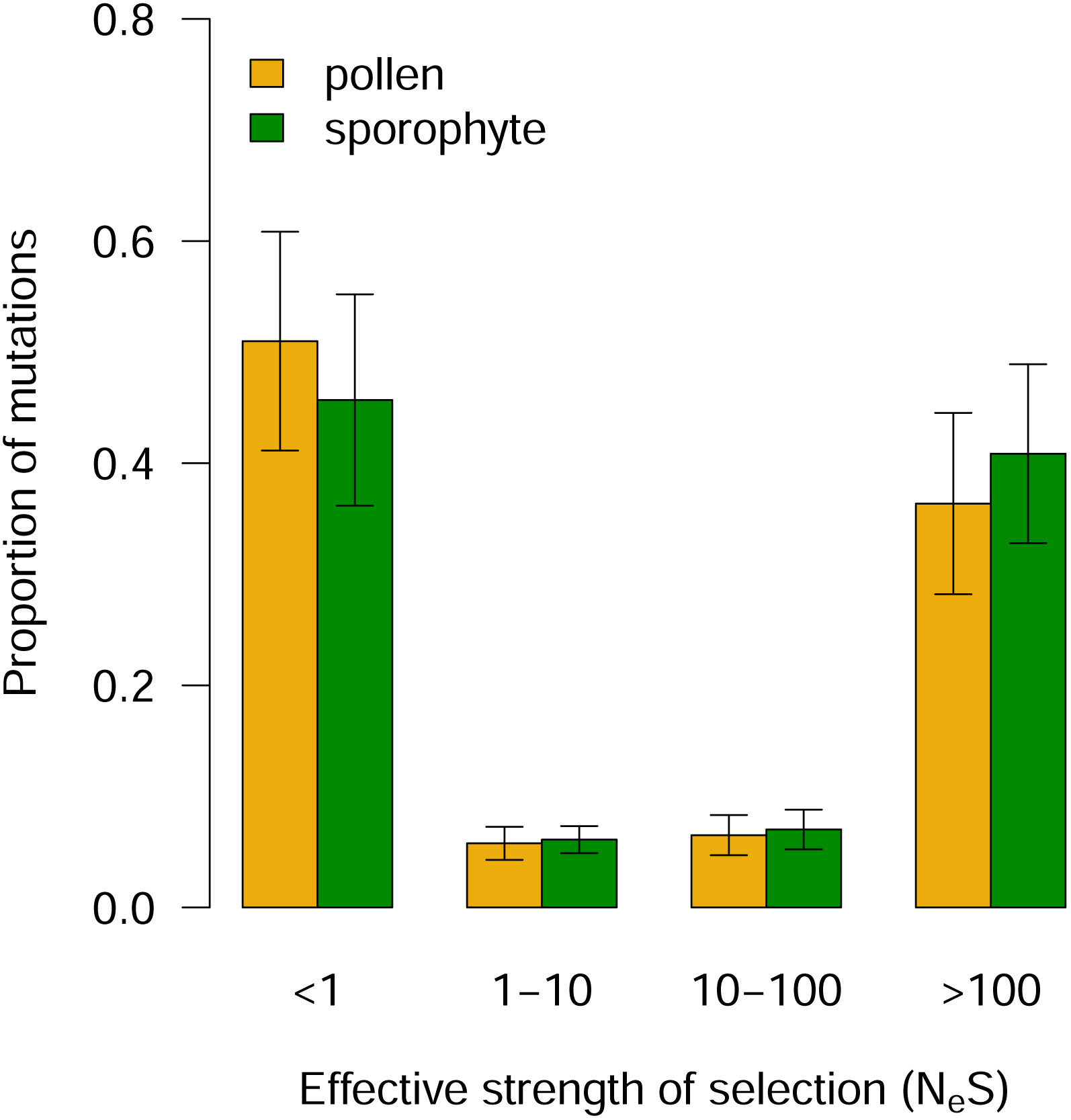
DFE for pollen and sporophyte-specific genes. Shown are the mean proportions of mutations in four N_*e*_s ranges with SDs

Using the same software we were unable to find evidence for positive selection for either group of genes, since *α* (proportion of sites under positive selection) was not significantly greater than zero in any of 10 random samples (mean: -1.6 in pollen; -1.9 in sporophyte genes). We also calculated *α* separately for each gene via an extension of the McDonald-Kreitman test presented in Smith and Eyre-Walker (2002). A slightly higher proportion (20.0%) of pollen genes had a positive value for *α* compared to 19.3% for sporophyte genes. A mean *α* of -3.5 in pollen genes and -3.9 in sporophyte genes, indicates, however, the prevalence of purifying selection. We conducted a further analysis to investigate levels of positive selection, which does not rely on polymorphism data. On a multi-sequence alignment containing single orthologues from each of the three species, *A. thaliana*, *A. lyrata* and *C. rubella*, we allowed dN/dS to vary among sites in order to detect sites under positive selection using codeml in PAML (Yang, 2007). This analysis, suggested a much higher proportion of pollen-specific genes contained sites under positive selection (15.2% at p *<* 0.05; 9.1 % at p *<* 0.01) compared to sporophyte-specific genes (9.3% p *<* 0.05; 4.8% at p *<* 0.01). As expected, dN/dS was significantly higher within the genes containing sites under positive selection compared to genes with no evidence for positive selection (median of 0.338 compared to 0.179 for pollen genes, p = 3.8 x 10^−21^; 0.228 compared to 0.154 in sporophyte genes, 3.9 x 10^−24^). It appears, therefore, that at least a part of the difference in dN/dS is caused by a higher rate of adaptive fixations in pollen genes.

### Pollen-specific genes are more polymorphic than sporophyte-specific genes

Pollen-specific genes were more polymorphic than sporophyte-specific genes with both non-synonymous nucleotide diversity (*π_n_*) and non-synonymous Watterson’s theta (*θ_n_*) significantly higher in pollen-specific genes (fig. 4). Both *π* and *θ* at synonymous sites did not differ between sporophyte-and pollen-specific genes (p = 0.18 & 0.58, respectively). Each of the six correlates of dN/dS listed above also correlated significantly with *π_n_* and *θ_n_* (all negatively except gene length; table 4). Five of the six variables (average intron length was not significant) explained 8.57% of variation in *π_n_* in a principal component regression. The first PC contributed most (3.11%). Four of the six factors (expression level, GC content, codon bias variance, and gene density) explained a total of 7.76% of the variation in *θ_n_* (first PC: 7.38%). For each model the first PC was implemented in an ANCOVA testing the influence of life-stage as a co-variate. *θ_n_* remained significantly higher for pollen-specific genes (p = 6.4 x 10^−61^; fig. 5(b)). PC1 had a significantly greater influence on *π_n_* for sporophyte genes (slope: -0.195) than on pollen genes (slope: -0.109; p = 7.2 x 10^−4^; Fig. 5(a)). We therefore tested the significance of difference in *π_n_*within 5 equal bins along the PC1 axis. In the 2nd to the 5th 20% quantiles *π_n_* was significantly higher within pollen genes, there was no difference in the first quantile (table 7).

**Figure 4:**
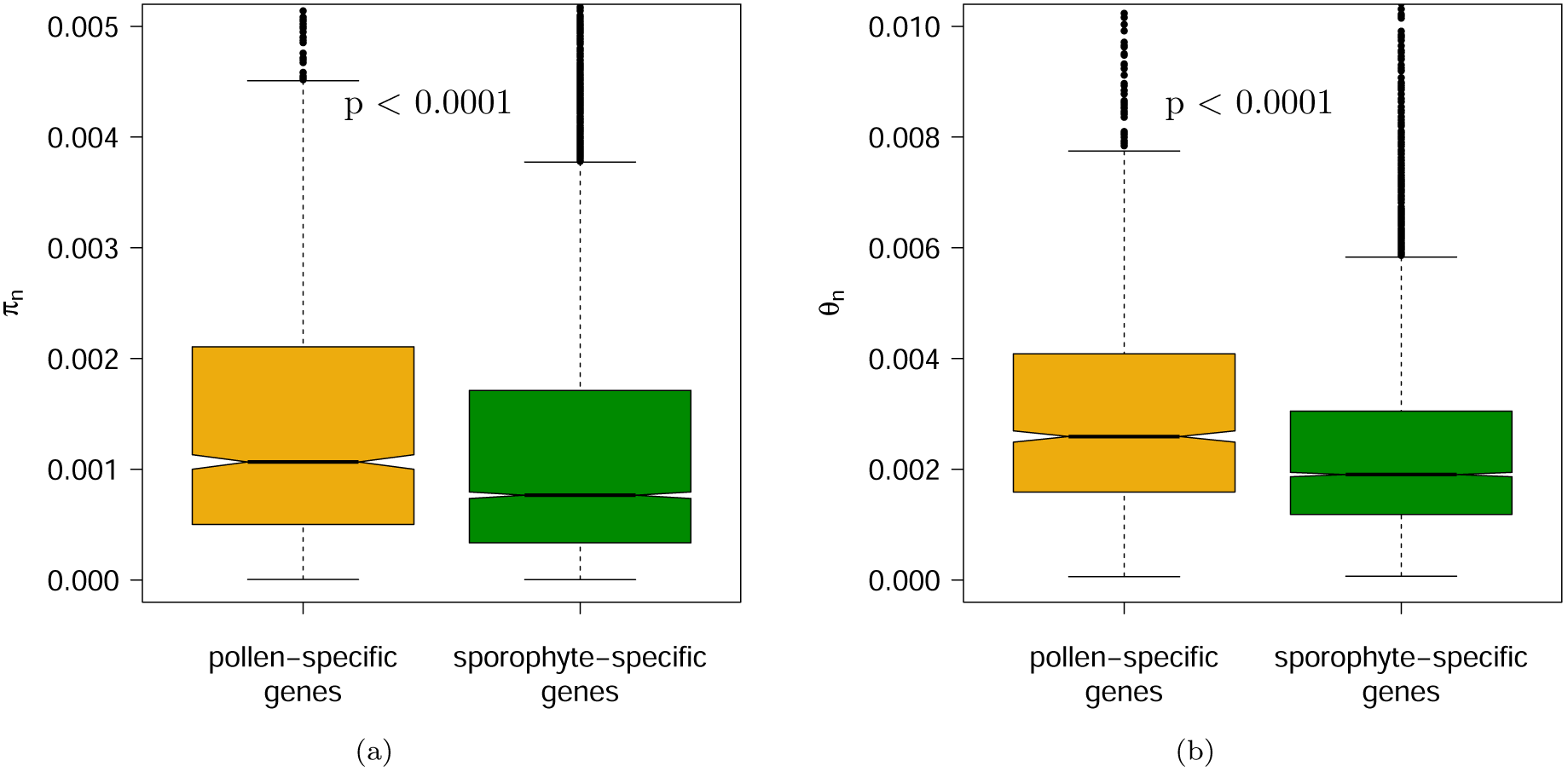
Non-synonymous nucleotide diversity (a) and non-synonymous Watterson’s theta (b) within pollen-specific and sporophyte-specific genes. Significance tested with Mann Whitney U test.

**Figure 5:**
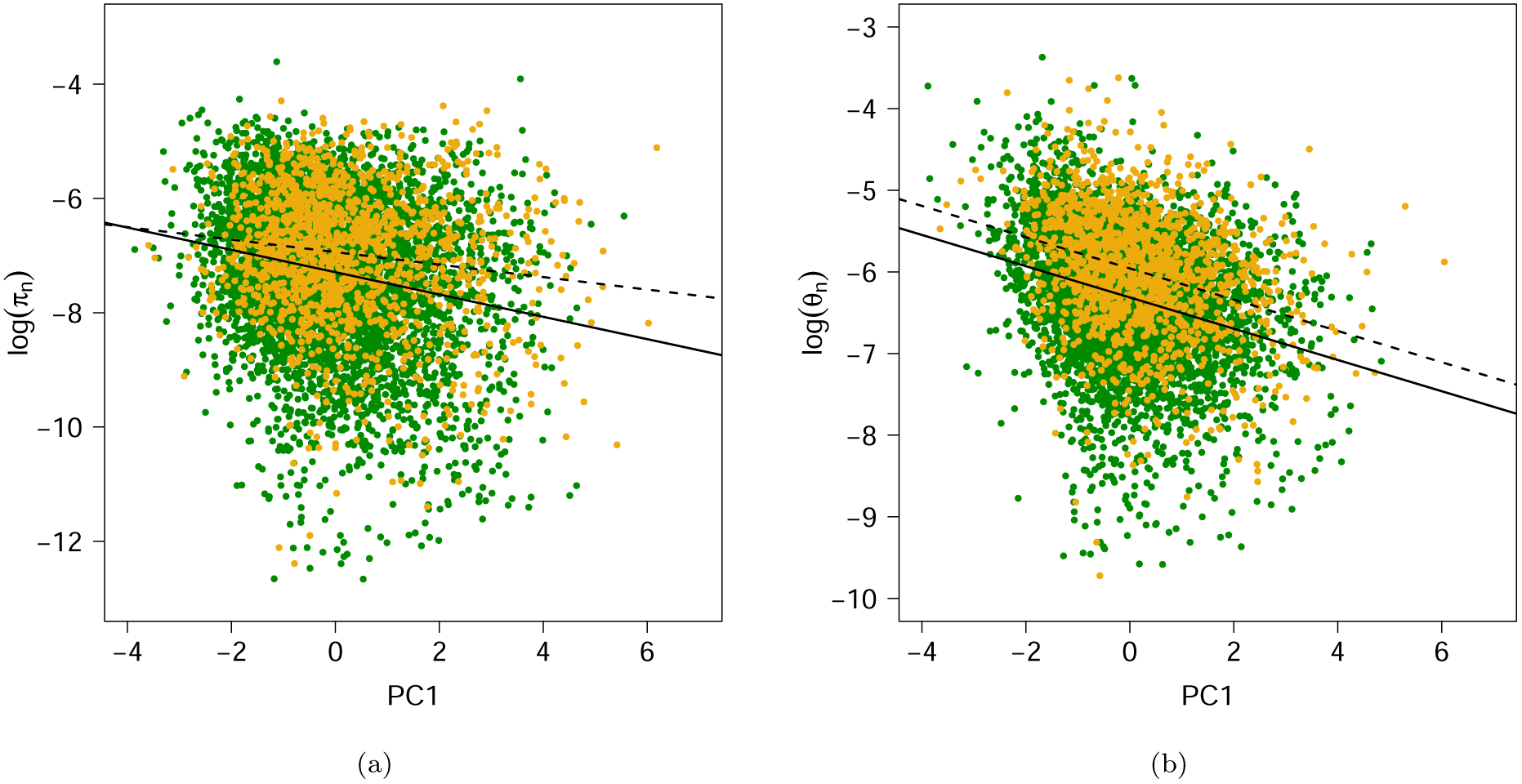
ANCOVA analysis with PC1 (6 genomic variables) as continuous variable reveals both higher *π_n_* and higher *θ_n_* (b) among pollen-specific (dark grey points and dashed line) than sporophyte-specific genes (light grey points and solid line).

**Table 7:** Nonsynonymous pi within 5 equal bins along the PC1 axis. Shown are medians (means).

|  | < 20% |  | 20% - 40% |  | 40% - 60% |  | 60% - 80% |  | > 80% |  |
| --- | --- | --- | --- | --- | --- | --- | --- | --- | --- | --- |
| Pollen | 1.0x10<sup>-3</sup> | (1.7x10<sup>-3</sup>) | 1.1x10<sup>-3</sup> | (1.7x10<sup>-3</sup>) | 1.1x10<sup>-3</sup> | (1.7x10<sup>-3</sup>) | 1.1x10<sup>-3</sup> | (1.6x10<sup>-3</sup>) | 8.4x10<sup>-4</sup> | (1.5x10<sup>-3</sup>) |
| Sporophyte | 1.0x10<sup>-3</sup> | (1.7x10<sup>-3</sup>) | 8.6x10<sup>-4</sup> | (1.4x10<sup>-3</sup>) | 7.2x10<sup>-4</sup> | (1.2x10<sup>-3</sup>) | 6.7x10<sup>-4</sup> | (1.1x10<sup>-3</sup>) | 6.0x10<sup>-4</sup> | (1.0x10<sup>-3</sup>) |
| p | ns |  | 1.1x10<sup>-3</sup> |  | 1.1x10<sup>-8</sup> |  | 5.8x10<sup>-6</sup> |  | 1.0x10<sup>-5</sup> |  |

### Higher frequency of deleterious mutations in pollen-specific genes

Higher polymorphism levels may indicate relaxed purifying selection on pollen-specific genes. To test this hypothesis further we investigated the frequency of putatively deleterious mutations - premature stop codons and frameshift mutations - within the 269 *A. thaliana* strains. Stop codon frequency, defined here as the relative number of unique alternative alleles due to premature stop codons occurring within the 269 strains, was significantly higher within pollen-specific genes (mean: 0.063 *±*0.004; sporophyte mean: 0.049 *±*0.002; p = 4.1 x 10^−15^; Mann Whitney U test; fig. 6). The frequency of strains containing at least one frameshift mutation was also significantly higher for pollen-specific genes (mean: 0.021 *±*0.002) compared to sporophyte-specific genes (mean: 0.014 *±*0.001; p = 6.6 x 10^−22^; fig. 6). Significant correlations existed between these measures of deleterious mutations and the six correlates of dN/dS (table 4).

**Figure 6:**
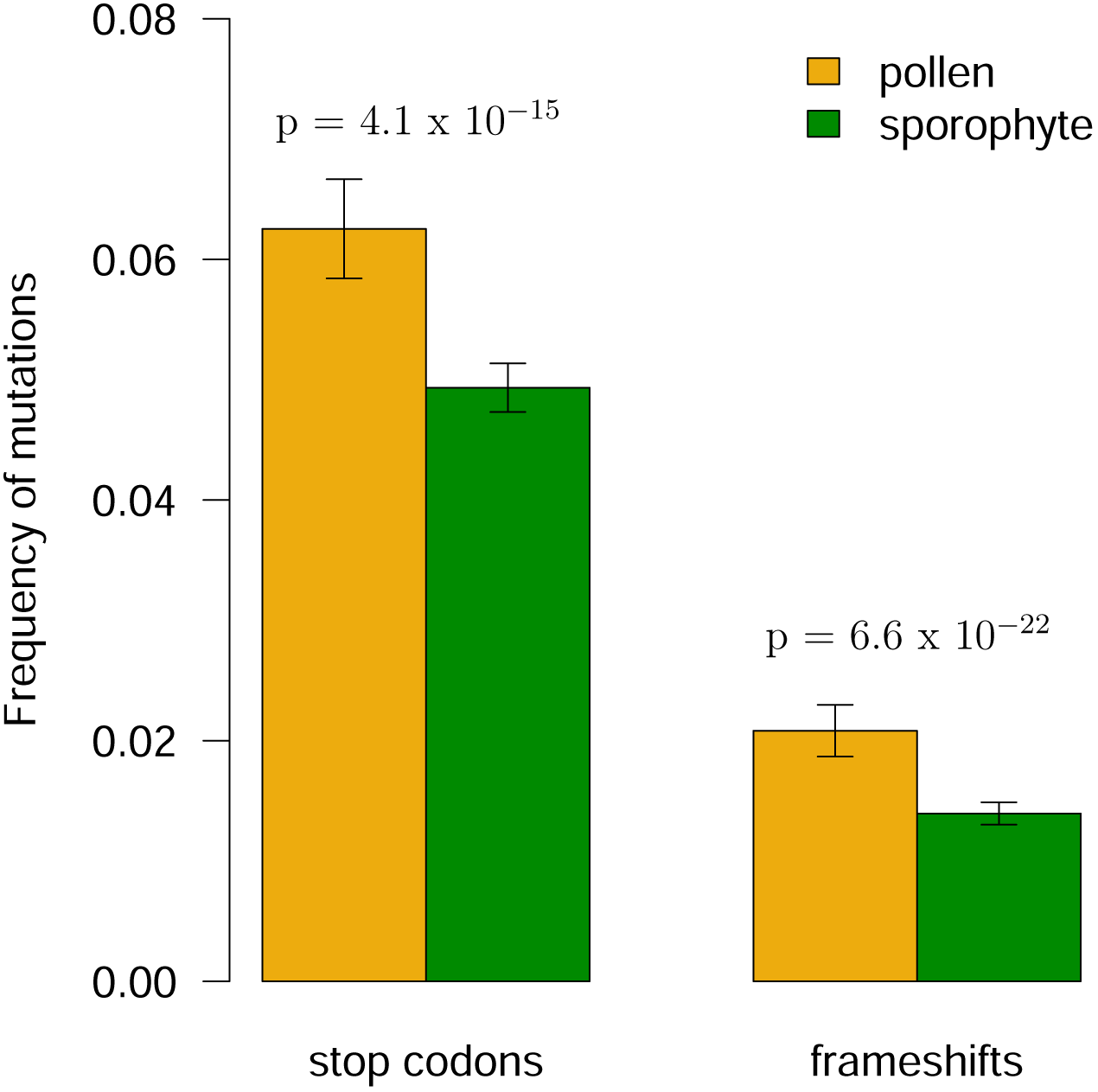
Frequency of alleles containing premature stop codon mutations and frameshift mutations in pollen-specific and sporophyte-specific genes. Significance tested with Mann Whitney U test.

In a principal component regression analysis all six predictors (expression level, codon bias variance, GC content, gene length, average intron length and gene density) were significantly correlated with stop codon frequency. The six predictors explained a total of 20.04% of the variation in stop codon frequency, 17.42% explained by the first PC. Within an ANCOVA with life-stage as the binary co-variant the frequency of premature stop codons remained higher within pollen-specific genes for the majority of PC1 (fig. 7(a)). The slopes differed significantly but the frequency of stop codons was significantly higher for pollen genes within the second to fifth 20% quantiles (table 8).

**Figure 7:**
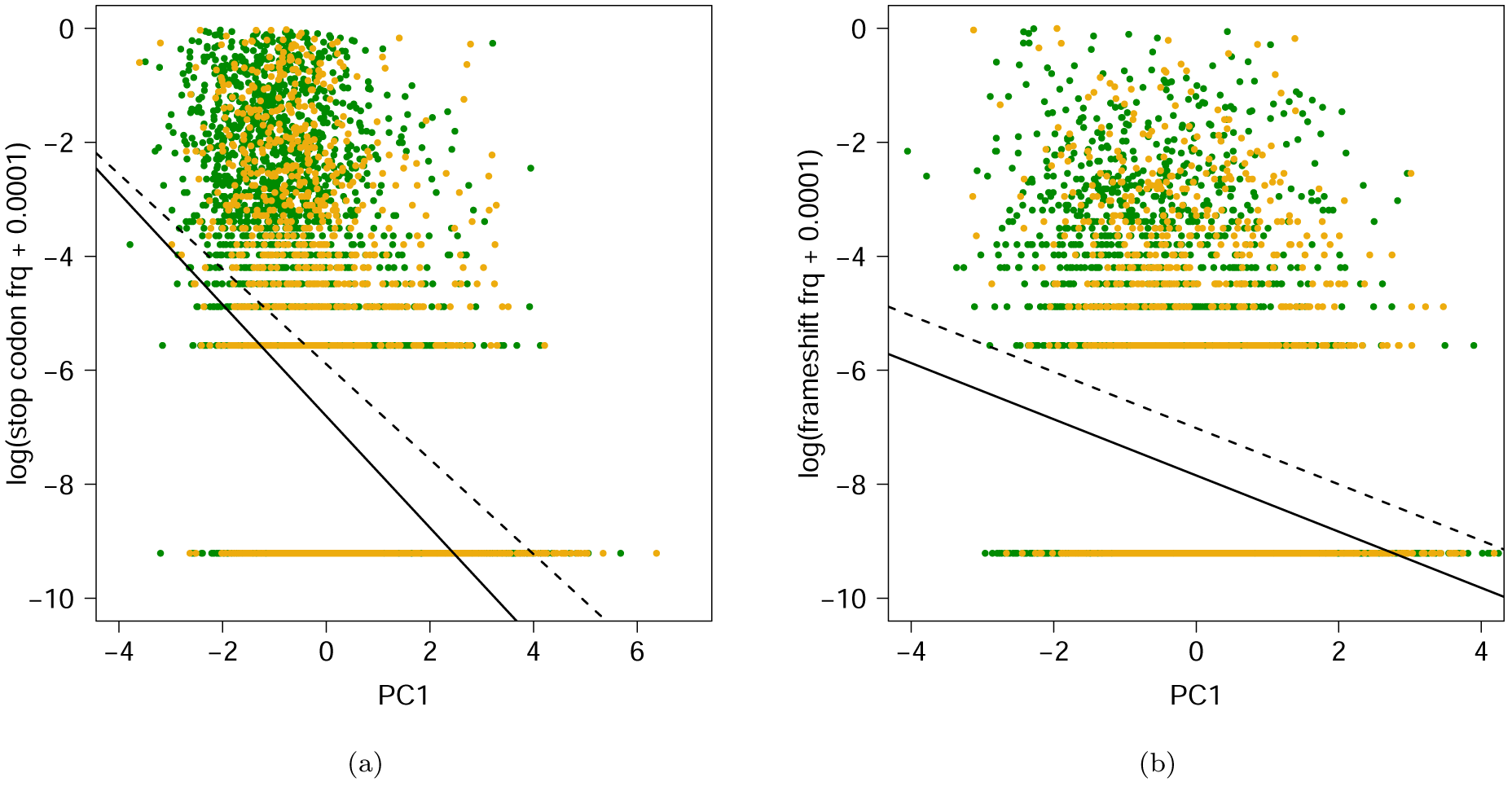
ANCOVA analysis with PC1 (6 genomic variables) as continuous variable reveals significantly higher frequency of stop codon mutations (a) and frameshift mutations (b) among pollen-specific (dark grey points and dashed line) than sporophyte-specific genes (light grey points and solid line).

**Table 8:**
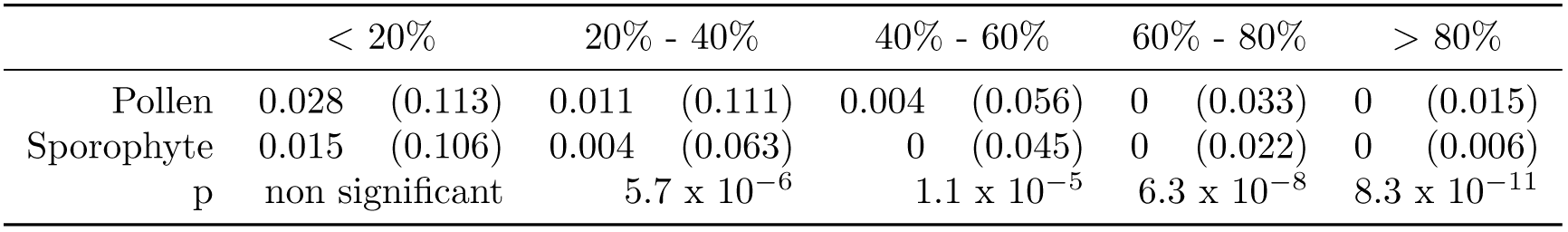
Frequency of stop codons within 5 equal bins along the PC1 axis. Shown are medians (means).

|  | < 20% |  | 20% - 40% |  | 40% - 60% |  | 60% - 80% |  | > 80% |  |
| --- | --- | --- | --- | --- | --- | --- | --- | --- | --- | --- |
| Pollen | 0.028 | (0.113) | 0.011 | (0.111) | 0.004 | (0.056) | 0 | (0.033) | 0 | (0.015) |
| Sporophyte | 0.015 | (0.106) | 0.004 | (0.063) | 0 | (0.045) | 0 | (0.022) | 0 | (0.006) |
| p | non significant |  | 5.7 x 10<sup>-6</sup> |  | 1.1 x 10<sup>-5</sup> |  | 6.3 x 10<sup>-8</sup> |  | 8.3 x 10<sup>-11</sup> |  |

**Table 9:** Differences in 6 genomic variables between pollen-specific genes and genes limited to one of three sporophytic tissues. Values are means ± standard error of the mean; significance was tested with Mann Whitney U test; p-values are Bonferroni corrected for multiple testing.

|  | Pollen-specific genes |  |  | guard cell, xylem or root hair |  |  | p |
| --- | --- | --- | --- | --- | --- | --- | --- |
| Expression level | 2,562.30 | ± 86.49 | > | 446.24 | ± 26.82 | | 1.0 x 10<sup>-77</sup> |
| GC content (%) | 44.20 | ± 0.08 | < | 44.80 | ± 0.17 | | 4.5 x 10<sup>-3</sup> |
| Codon bias variance | 0.46 | ± 0.01 | > | 0.39 | ± 0.01 | | 2.2 x 10<sup>-6</sup> |
| gene length | 1,570.30 | ± 24.41 | = | 1,561.71 | ± 36.20 | | not significant |
| average intron length | 124.44 | ± 3.23 | = | 152.49 | ± 9.14 | | not significant |
| gene density (per 100kb) | 29.99 | ± 0.12 | = | 29.48 | ± 0.30 | | not significant |

**Table 10:** Expression data sets.

|  | Dataset | Description | Chips | Original source |
| --- | --- | --- | --- | --- |
| Haploid | UNM | Uninucleate microspore | 2 | Honys & Twell, 2004 |
|  | BCP | Bicellular pollen | 2 | Honys & Twell, 2004 |
|  | TCP | Tricellular Pollen | 2 | Honys & Twell, 2004 |
|  | MPG | Mature Pollen | 2 | Honys & Twell, 2004 |
|  | GP* | Pollen Tube Grouped | 6 | Qin et al., 2009 ; Wang et al., 2008 |
|  | PT4* | Pollen Tube Grouped | 6 | Qin et al., 2009 ; Wang et al., 2008 |
|  | SPC | Sperm Cell | 3 | Borges et al., 2008 |
| Diploid | SL | Silique | 30 | NASC |
|  | LF | Leaves | 36 | NASC |
|  | GC | Guard Cell | 3 | NASC |
|  | PT** | Petiole | 3 | NASC |
|  | ST | Stems | 2 | NASC |
|  | HP | Hypocotyl | 8 | NASC |
|  | XL | Xylem | 3 | NASC |
|  | CR | Cork | 3 | NASC |
|  | RT | Roots | 11 | NASC |
|  | RH | Root hair elongation zone | 3 | NASC |

NASC: Nottingham Arabidopsis Stock Centre.
\* GP and PT4 were combined to one data set called PT, selecting the highest expression level of the two for each gene.
\*\* Renamed PET

Four of the predictors (expression level, GC content, gene length and gene density) were also significantly correlated with the frequency of frameshift mutations. However, the four variables only explained a total of 5.49% of variation (first PC 5.08%). In an ANCOVA analysis frameshift mutations remained significantly more frequent within pollen-specific genes when controlling for the predictors via the first PC (fig. 7(b)).

### Tissue specific genes

Tissue specificity has been shown to be negatively correlated with selection efficiency (Duret and Mouchiroud, 2000; Liao *et al.*, 2006; Slotte *et al.*, 2011). The on average greater tissue specificity in pollen-specific genes compared to sporophyte specific genes could therefore potentially explain the higher polymorphism levels and higher frequency of deleterious mutations found in pollen-specific genes. In order to control for this potential bias we compared dN/dS, polymorphism levels and the frequency of deleterious alleles in pollen-specific genes with a group of 340 genes with expression limited to a single sporophyte cell type (guard cell, xylem or root hair). To further test for the effect of tissue specificity, these groups were also compared against 2543 genes which were expressed in at least 5 sporophytic tissues.

In this tissue-specificity controlled comparison, dN/dS did not differ between pollen-specific and the tissue-specific sporophyte gene set. However, dN/dS was significantly higher in pollen-specific genes (p = 1.7 x 10^−27^) and tissue specific sporophyte genes (p = 1.0 x 10^−9^; fig. 8) compared to broadly expressed sporophyte-specific genes. In a principal components regression only expression level and GC content had a significant effect on dN/dS, explaining 8.63% of variation. The PC1 (8.60%) was then mapped against dN/dS in an ANCOVA on the two levels pollen-specific genes and tissue-specific, sporophytic genes. dN/dS was significantly higher for pollen genes than tissue-specific, sporophytic genes when controlling for PC1 (p = 1.4 x 10^−3^; figure 9).

**Figure 8:**
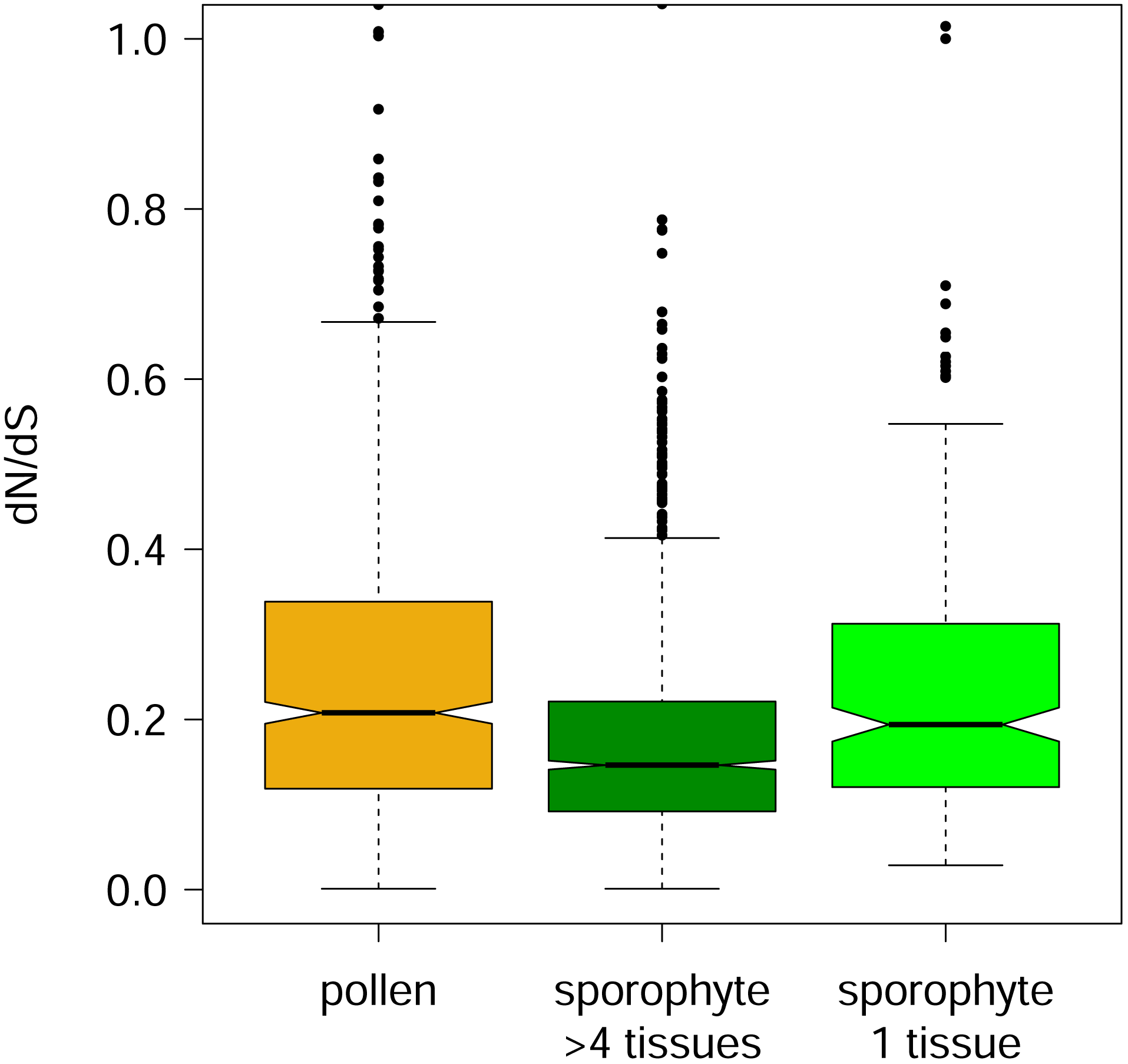
dN/dS within pollen-specific genes, broadly expressed sporophytic genes (at least 5 tissues) and tissue specific genes (expression restricted to guard cell, xylem or root hair tissues).

**Figure 9:**
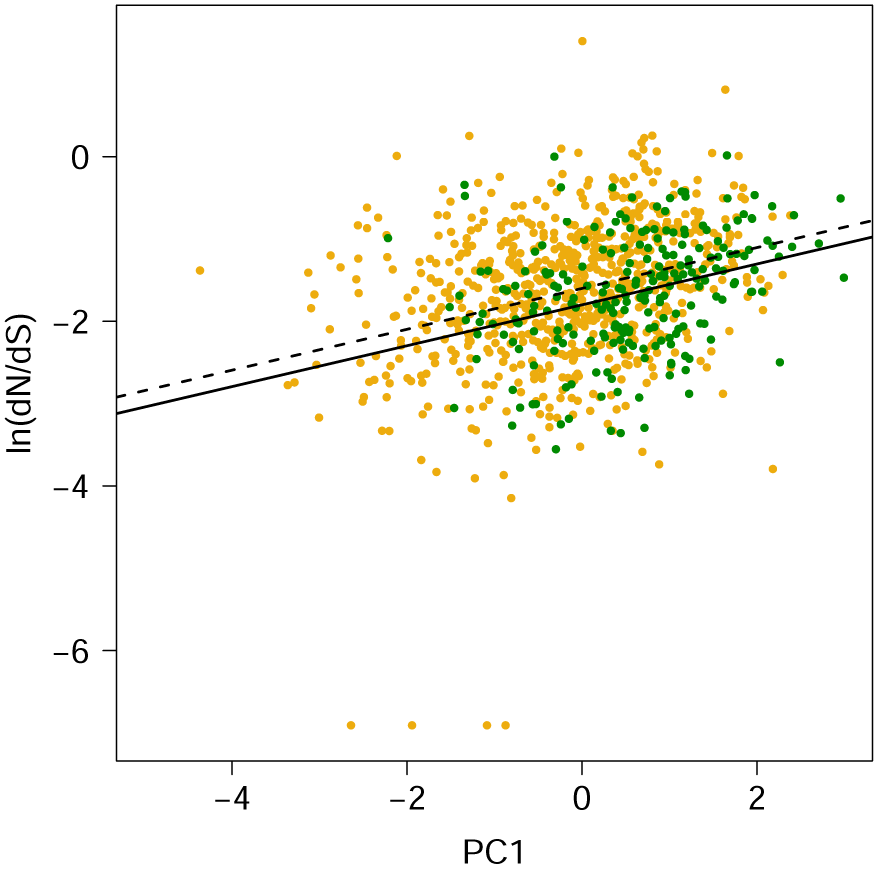
ANCOVA analysis of dN/dS within pollen-specific (yellow points and dashed line) and tissue specific, sporophyte genes (green points and solid line) with PC1 (expression and GC content) as the continuous variable.

Similarly, *π_n_* and *θ_n_* did not differ between pollen-specific and the tissue-specific sporophyte gene set. However, they were both significantly higher in pollen-specific genes (p = 1.6 x 10^−30^ & 8.4 x 10^−75^, respectively) and tissue specific sporophyte genes (p = 7.1 x 10^−13^ & 2.7 x 10^−26^; fig. 10) compared to broadly expressed sporophyte-specific genes. In a principal components regression, expression level and GC content had a significant effect on *π_n_*, explaining 5.30% of variation. The first PC (5.06%) was plotted against *π_n_* in an ANCOVA within pollen-specific and tissue-specific sporophytic genes. *π_n_* was significantly higher for pollen-specific genes compared to tissue-specific, sporophytic genes when controlling for PC1 (p = 6.5 x 10^−3^; figure 11(a)). In a similar analysis for *θ_n_*, all 6 parameters (expression level, GC content, codon bias variance, gene length, average intron length and gene density) significantly contributed to 18.82% of variation. The second PC was largest (9.55%) and was plotted against *θ_n_* in an ANCOVA (figure 11(b)). When controlling for PC2 *θ_n_* was significantly higher within pollen-specific genes (p = 2.9 x 10^−7^) compared to tissue-specific, sporophytic genes.

**Figure 10:**
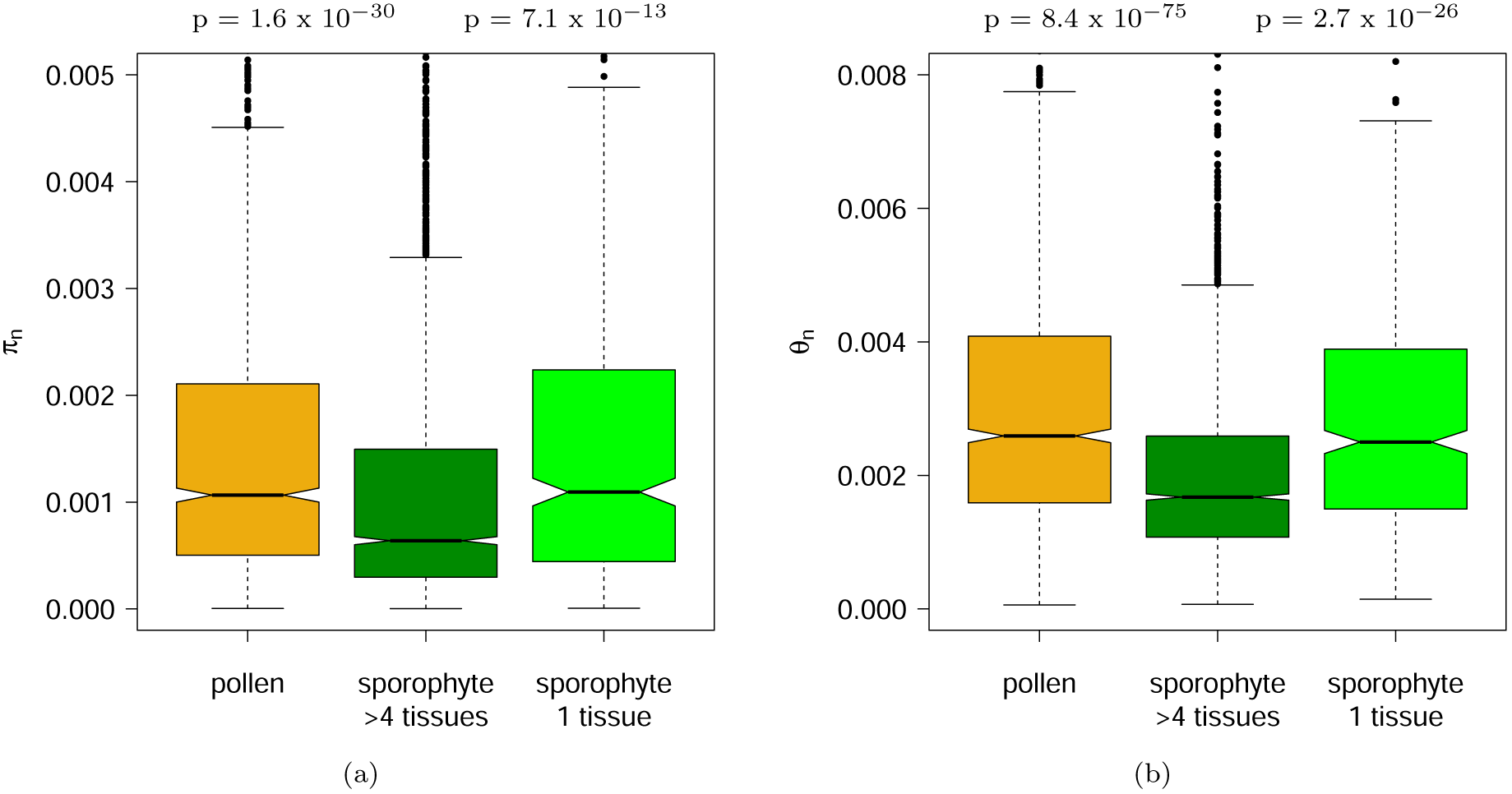
Non-synonymous nucleotide diversity (a) and non-synonymous Watterson’s theta (b) within pollen-specific genes, broadly expressed sporophyte-specific genes and genes specific to guard cells, xylem or root hair.

**Figure 11:**
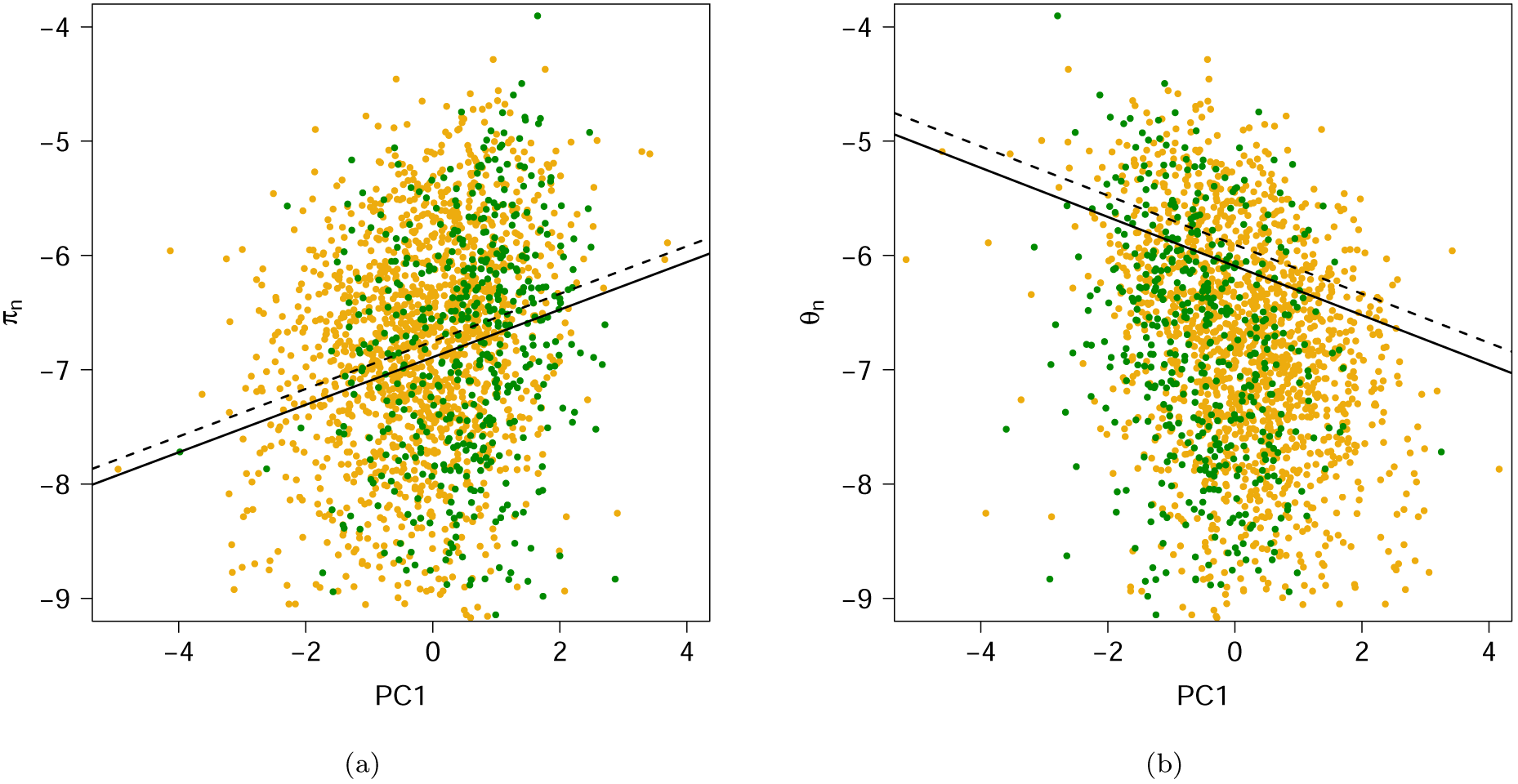
ANCOVAS comparing *π_n_* (a) and *θ_n_* (b) within pollen-limited genes (yellow points and dashed line) to tissue-specific, sporophytic genes (green points and solid line) while controlling for the first PC of a PCR.

Premature stop codons remained significantly more frequent in pollen-specific genes than in sporophytic, tissue specific genes (p = 0.033), and broadly expressed, sporophytic genes (p = 3.0 x 10^−14^; fig. 12). Premature stop codons were more frequent in tissue specific, sporophytic tissues (mean 0.057 *±* x 10^−3^) compared to broadly expressed sporophytic genes (0.051 *±* 3.2 x 10^−3^) but not significantly. There was no significant difference in the frequency of frameshift mutations between pollen-specific genes and tissue-specific, sporophytic genes but the frequency was significantly higher in both groups compared to broadly expressed, sporophytic genes (p = 2.0 x 10^−34^ & 1.7 x 10^−14^; figure 12). GC content, codon bias variance, gene length, average intron length and gene density had a significant effect on 19.03% variation in the frequency of stop codons. The first PC was largest (16.44%) and was implemented in an ANCOVA as the continuous variable (fig. 13(a)). Due to a significant interaction between the two groups, pollen-specific genes and tissue-specific, sporophytic genes, differences in stop codon frequencies were measured within 5 equal bins along the PC1 axis. The frequency of stop codon mutations did not differ significantly within the first four quantiles but was significantly higher within pollen-specific genes in the fifth quantile (PC1 *>* 1.14; p = 5.7 x 10^−5^). The analysis was repeated for the frequency of frameshift mutations. Expression level, GC content, codon bias variance and gene length explained a total of 6.35% variation. PC2 was largest with 3.24% so was implemented in an ANCOVA. The frequency of frameshift mutations was significantly higher within pollen-specific genes compared to tissue-specific, sporophytic genes when controlling for PC2 (p = 0.017; figure 13(b)).

**Figure 12:**
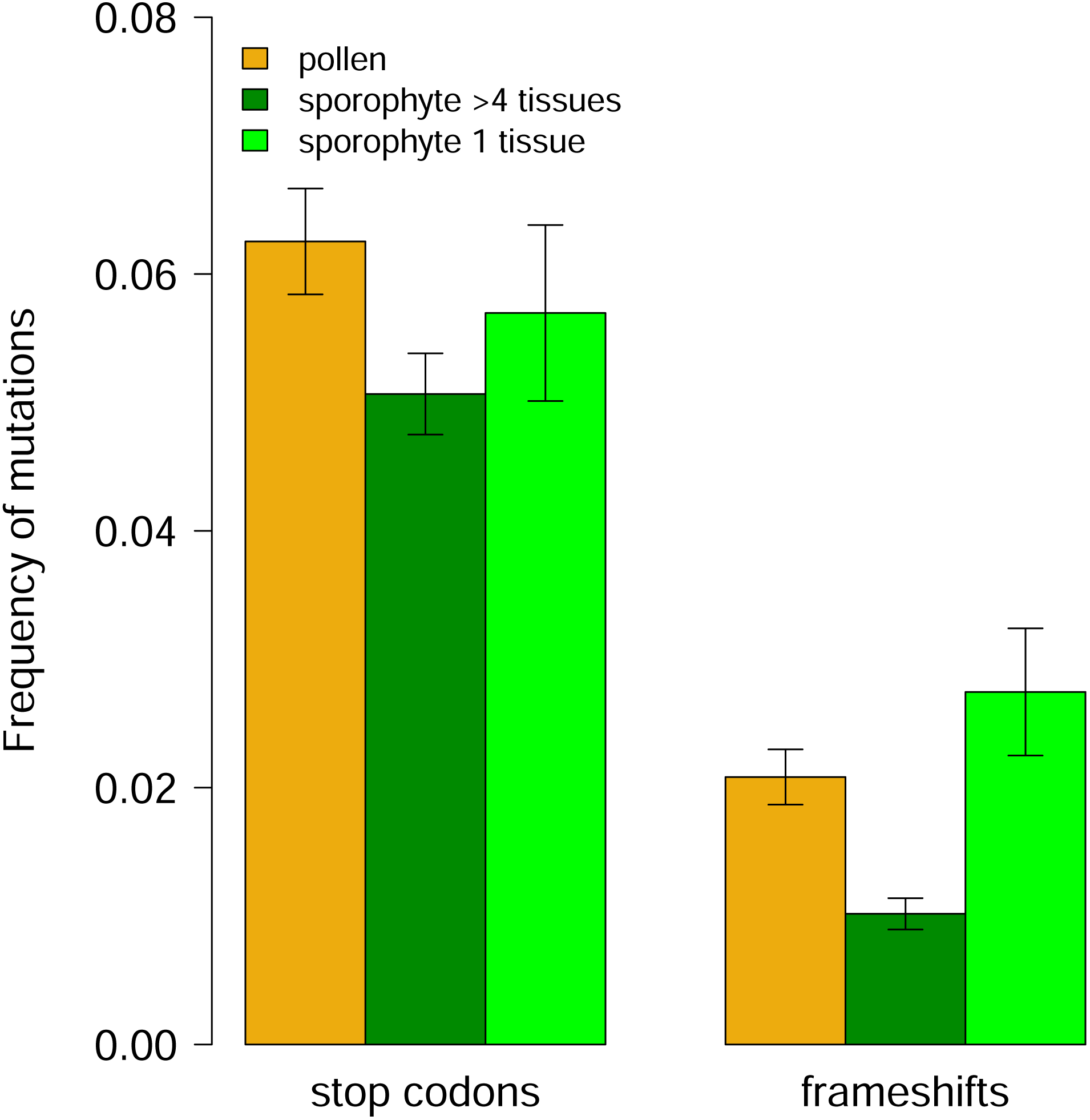
Frequency of stop codon and frameshift mutations within pollen-specific genes, broadly expressed sporophytic genes (at least 5 tissues) and tissue specific genes (expression restricted to guard cell, xylem or root hair tissues).

**Figure 13:**
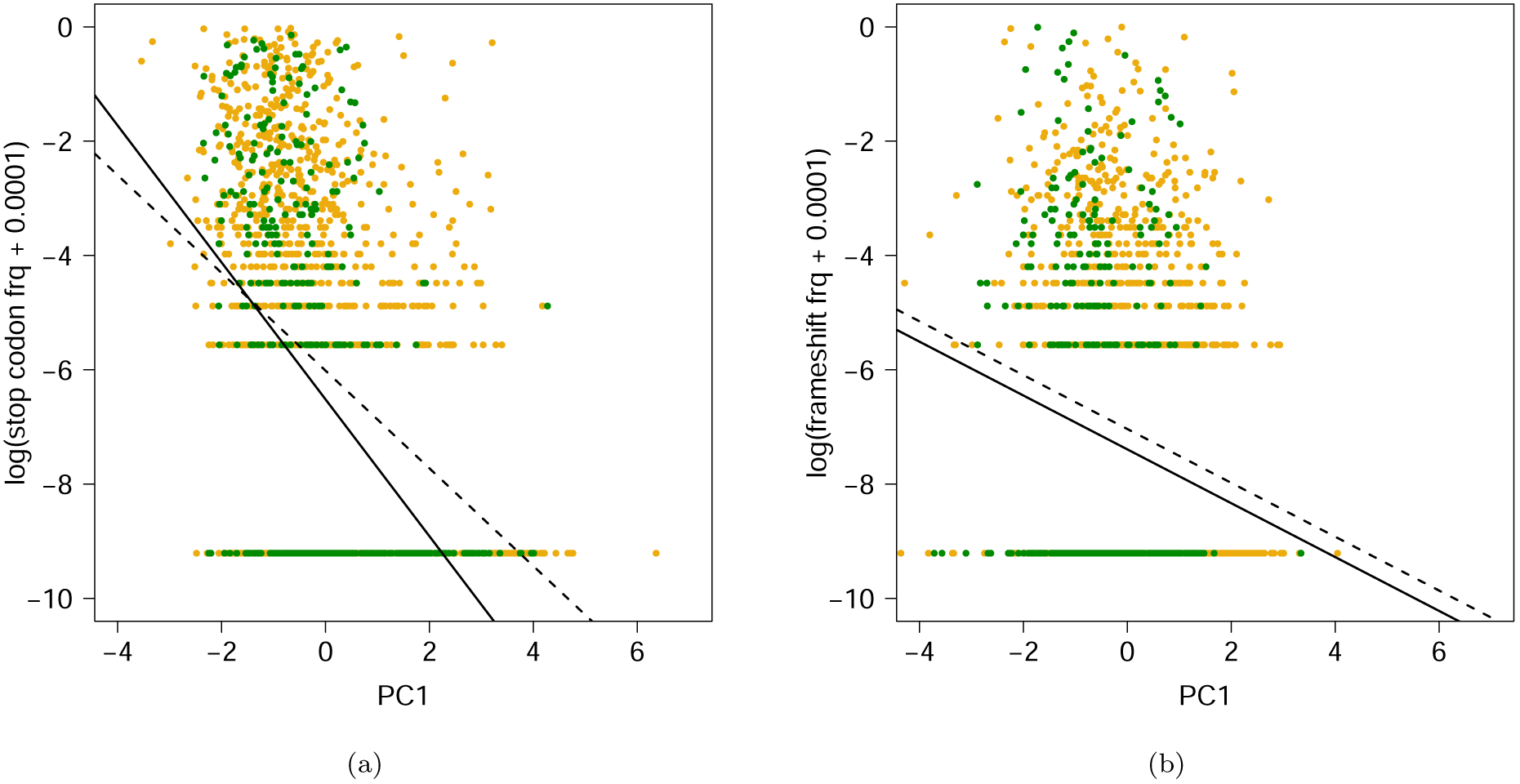
ANCOVAS comparing the frequency of stop codon mutations (a) and frameshift mutations (a) within pollen-limited genes (yellow points and dashed line) to tissue-specific, sporophytic genes (green points and solid line) while controlling for the first PC of a PCR.

## Discussion

Our analysis showed that protein divergence, polymorphism levels and the frequency of deleterious mutations were significantly higher within pollen-specific genes compared to sporophyte-specific genes. These differences remained when controlling for expression level, GC content, codon bias variance, gene length, average intron length and gene density.

### Evolutionary rates higher within pollen-specific genes

Protein divergence rates (dN/dS) were on average 37% higher in pollen-specific genes compared to sporophyte-specific genes. This is comparable to the findings presented by Szövényi *et al.* (2013), who found dN/dS to be 39% or 81% higher in pollen genes for *A. thaliana* depending on the data set. In a further paper, no difference in dN/dS could be found between pollen-specific and non-reproductive genes for *A. thaliana* (Gossmann *et al.*, 2013). This discrepancy was most likely caused by the method of gene selection. In the Szövényi *et al.* (2013) study, as in the current study, genes with exclusive expression within sporophytic or pollen tissues were analysed. In the Gossmann *et al.* (2013) paper, on the other hand, genes were selected more inclusively, labelling a gene as pollen-enriched if expression was significantly higher at a fold change greater than 4 within different comparisons. This means that at least some of the sporophyte genes discussed in the Gossmann *et al.* (2013) study will also be expressed to some extent within pollen tissues and are therefore exposed to haploid selection. Even a low level of expression in haploid tissues may be sufficient to counteract the effect of masking, which would explain the lack of difference in evolutionary rates detected. It appears then that the genes, which are exclusively expressed in pollen or sporophytic tissues, may be causing the significantly different dN/dS rates we observe here.

These higher dN/dS values can, in part, be explained by stronger positive selection acting on pollen-specific genes compared to sporophyte specific genes, as indicated by a greater proportion of pollen-specific genes containing sites under positive selection (15.2% compared to 9.3%). However, an analysis of the distribution of fitness effects of new nonsynonymous mutations revealed a higher frequency of effectively neutral mutations within pollen-specific genes. This indicates purifying selection is more relaxed within pollen-specific genes, suggesting the higher dN/dS rate within pollen-specific genes may have been caused by a greater proportion of slightly deleterious substitutions due to random drift.

### Polymorphism levels suggest relaxed selection on pollen-specific genes

Polymorphism levels were significantly higher within pollen-specific genes. Both Watterson’s *θ* and *π* of non-synonymous sites remained significantly higher within pollen-specific genes when controlling for expression and five further genomic differences (GC content, codon bias variance, gene length, average intron length and gene density). In one of two recent studies, higher polymorphism rates were also found in pollen-specific genes for *A. thaliana* (Szövényi *et al.*, 2013). In the second study, however, no difference was found between pollen-specific genes in general and random, non-reproductive genes in terms of nucleotide diversity (Gossmann *et al.*, 2013), which, as discussed in the previous section, is possibly due to the more inclusive choice of genes in that study.

We also found significantly higher levels of putatively deleterious alleles (premature stop codons and frameshift mutations) within pollen-specific genes. This supports the conclusions of Szövényi *et al.* (2013) that the raised polymorphism levels indicate relaxed purifying selection on pollen-specific genes. In other words, comparatively weaker selective constraints are allowing deleterious alleles to accumulate at a greater rate within pollen-specific genes compared to those whose expression is restricted to the sporophyte.

### Has there been a recent shift in selection strength?

The patterns in our data are compatible with a change in selection efficacy that is likely to have taken place since the speciation of *A. thaliana* and *A. lyrata*. The relatively recent switch from self-incompatibility to self-compatability in *A. thaliana* (ca. 1MYA; Tang *et al.* 2007) explains why we have observed evidence for relaxed selection in polymorphism levels but stronger positive selection in divergence data for pollen-specific genes. The divergence data used to calculate dN/dS mainly represent a prolonged period of outcrossing (~ 12 MYA), since the speciation of *A. thaliana* from *A. lyrata* occurred roughly 13 million years ago (Beilstein *et al.*, 2010). In contrast, the polymorphism data and frequencies of putative deleterious alleles reflect the recent selective effects of high selfing rates. This may also explain why slightly more relaxed purifying selection was discovered for pollen-specific genes by the DoFE analysis, since it also relies on polymoprhism data.

The evidence we have found for a more recent weaker selection on pollen-specific genes contrasts with findings for the outcrossing *Capsella grandiflora* (Arunkumar *et al.*, 2013). In that study the more efficient purifying and adaptive selection on pollen genes was linked to two possible factors: haploid expression and pollen competition. *A. thaliana* is a highly self-fertilizing species with selfing rates generally in the range of 95 - 99% (Platt *et al.*, 2010), so haploid expression is unlikely to improve the efficacy of selection on pollen-specific genes relative to sporophyte genes. This is because most individuals found in natural populations are homozygous for the majority of loci, reducing the likelihood that deleterious alleles are masked in heterozygous state when expressed in a diploid tissue (Platt *et al.*, 2010). A reduction in pollen competition can also be expected due to the probably limited number of pollen genotypes in highly selfing populations (Charlesworth and Charlesworth, 1992; Mazer *et al.*, 2010). However, outcrossing does occur in natural *A. thaliana* populations with one study reporting an effective outcrossing rate in one German population of 14.5% (Bomblies *et al.*, 2010). Nevertheless, it appears that these generally rare outcrossing events may not be sufficient to prevent a reduction in pollen competition for *A. thaliana*.

So if we assume both masking and pollen competition are negligible forces when comparing selection on pollen-specific genes to sporophyte-specific genes, why is selection more relaxed on pollen-specific genes than sporophyte-specific genes rather than similar?

We have shown here that tissue specificity partly explains why selection is more relaxed on pollen genes. The full set of sporophyte-specific genes contains genes expressed across several tissues, and broadly expressed genes have been known to be under more efficient selection than tissue-specific genes due to their exposure to a higher number of selective constraints (Duret and Mouchiroud, 2000; Liao *et al.*, 2006; Slotte *et al.*, 2011). Both pollen-specific genes and genes limited to one of three sporophytic tissues (xylem, guard cell or root hair) showed raised levels of dN/dS, polymorphism and frequency of deleterious mutations compared to broadly expressed sporophyte-specific genes (expressed in at least 5 tissues). Tissue specificity appeared to explain, to a certain extent, the reduced selection efficacy in pollen-specific genes as there was no longer a significant difference in polymorphism levels (*θ_n_* and *π_n_*) or the frequency of frameshift mutations in pollen-specific genes compared to the tissue specific, sporophytic genes (the frequency of stop codon mutations remained significantly higher). However, tissue specificity alone only partly explains the apparent, current more relaxed selection on pollen-specific genes. Once further genomic features (expression level, GC content, codon bias variance, gene length, average intron length, gene density) were controlled for, all measures remained higher in pollen-specific genes even when compared to genes restricted to only one sporophytic tissue except for stop codon frequency.

The difference in rates of protein evolution between pollen-specific and sporophyte-specific genes could also be explained to a large extent by tissue specificity. As with polymorphism level and frequencies of deleterious mutations, dN/dS did not differ between pollen-specific and tissue specific, sporophytic genes. However, dN/dS was significantly higher in both gene groups compared to broadly expressed, sporophytic genes. This suggests that the specificity of pollen genes to a small set of tissues is responsible for their elevated rates of protein evolution rather than their specific association with pollen tissues. Again, this only partially explains the raised dN/dS levels, as when controlling for differences in expression level and GC content, dN/dS remained significantly higher within pollen-specific genes.

Previously reported similar findings indicating relaxed purifying selection in pollen specific genes in *thaliana* (Szövényi *et al.*, 2013) were explained as possibly resulting from a combination of high tissue specificity and higher expression noise in pollen compared to sporophytic genes. However, the authors did not compare selection on pollen genes to tissue specific sporophyte genes to isolate the effect of tissue specificity. We have shown here that tissue specificity does appear to play a role but does not alone explain the difference in selection strength between both groups of genes. Higher expression noise could then be an important factor influencing the level of deleterious alleles which exist for pollen genes in *A. thaliana*.

Expression noise has been found to reduce the efficacy of selection substantially and is expected to be considerably higher for haploid expressed genes (Wang and Zhang, 2011). It is therefore likely that in the absence of pollen competition and the masking of deleterious sporophyte-specific genes, expression noise and high tissue specificity become dominant factors for pollen-specific genes of selfing plants. The loss of self-incompatibility in *A. thaliana* may therefore have led to a reduction in selection efficacy and the accumulation of deleterious alleles in pollen-specific genes.

### Conclusion

We have shown that, as in many other taxa, genes expressed in male reproductive tissues evolve at a quicker rate than somatic genes in *A. thaliana*. The greater divergence of pollen proteins to both *A. lyrata* and *C. rubella* compared to sporophytic genes can be attributed to stronger positive and purifying selection. However, intra-specific polymorphism data indicate a strong shift in this selection pattern may have occurred. Since the more recent loss of incompatibility in *A. thaliana* selection appears to have become more relaxed in pollen-specific genes. This is likely due to a reduction in pollen competition and the masking of diploid, sporophytic genes as a result of high homozygosity levels. In outcrossing plants, haploid expression and pollen competition outweigh the negative impact of high tissue specificity and expression noise on the selection efficacy of pollen-specific genes. In the self-compatible *A. thaliana* high homozygosity has likely reduced the counteracting effects of pollen competition and haploid expression, leading to lower selection efficacy and an increased accumulation of deleterious mutations in pollen-specific compared to sporophyte-specific genes.

## Acknowledgements

MCH was supported by a PhD research grant from the Natural Environment Research Council (NERC). DT would like to acknowledge financial support from the UK Biotechnology and Biological Science Research Council (BBSRC).

## Author contributions

All four authors developed the project idea and were involved in the interpretation of data and finalization of the manuscript. MCH analyzed the data and drafted the manuscript

